# AutoTA: Galaxy Workflows for Reproducible and Automated Taxonomic Analysis using Qiime2

**DOI:** 10.1101/2024.04.29.591690

**Authors:** Atharva Tikhe, Shweta Jangam, Preeti Arora, Sanjay Gupte, Sarjan Shah

## Abstract

Metagenomic sequencing allows systematic characterization of microbial populations isolated from various environments of interest by bypassing the culturing of the isolates. Concomitant to advancement in sequencing techniques, analysis methods and softwares have also grown to be sophisticated and efficient. Qiime2 is a collection of python scripts which enables end-to-end analysis of metagenomic data. However, usage of latest and more complex databases for classification is hindered by requirement of high compute power. To aid cloud-based analysis, we present workflows for diversity analysis and taxonomic assignment which are the two most common and initial steps in a metagenomics experiments. The workflows are made in Galaxy which makes testing and analysing multiple datasets faster, in parallel, reproducible and flexible. The workflows can be integrated into a local Galaxy instance or used completely on the web which is of great importance to non-bioinformaticians and bench scientists.

## 1 Introduction

Metagenomics involves genome sequencing of complex and diverse microbial communities sampled from the environment. It is known that around 99% of microorganisms cannot be easily grown in a lab and hence the determination and analysis of actual diversity of microorganisms is limited in culture-dependent methodologies[1]. Metagenomic approaches bypass the culturing stage and directly sequence the samples isolated from various environments like ocean, soil and human body attempting to give a clearer picture of how microbial communities behave and how the community structure change in response to varying external factors. Generally, two of the most common methods in acquiring microbiome sequencing is either complete shotgun sequencing or more popular amplicon based sequencing targeting the 16S rRNA of bacteria. [2] Apart from 16S, other metabarcoding experiments are also performed viz. 18S or fungal ITS amplicons.

Bacterial 16S rRNA gene contains a hypervariable region in the range 30-100 bp and classified into regions V1-V9. The variability in these regions can be used as a standard for identification and taxonomic assignment of the organism [3]. There exist special tools for analyze the data generated from such amplicon-based sequencing experiments. The tool we focus on is qiime2 [4]. Qiime2 (which stands for ‘quantitative insights into microbial ecology’) is a software that provides end-to-end analysis capabilities through it’s vast collection of python scripts implemented in a plugin-based architecture. Adopting plugin-based architecture makes it easier for bioinformaticians to develop and integrate newer features in the form of python wrappers. However, hurdles in analyzing metagenomic data (shotgun or amplicon) arise, which are, (1) the requirement of powerful CPUs and high order of RAM for example, training taxonomic classifiers, which are usually deployed as clusters or high-end servers. (2) requirement of basic knowledge of linux environments, software installation and bioinformatics expertise in handling and manipulating data. Although these tools are widely used to analyse large data in metagenomic experiments, they still reqiure some level of familiarity with the command line and linux environment and as a result bench scientists with limited bioinformatics experience find it difficult to setup and run pipelines like qiime2 even in a docker container.

Use of web/Graphical User Interface (GUI) applications, like the Galaxy project [5] or the Genomics Virtual Lab [6] have gained attention of the scientific communities mainly because they bypass the need of setting up all the tools and using the command line and additionally, absense of in-house high-end computational infrastructure. These applications have pre-installed and pre-configured tools and do not require significant resources on the client side as the major analytical processing occurs on the server side. The Galaxy platform allows reproducible and scalable NGS data analysis by providing scientists with an intuitive GUI. It enables the scientist to rapidly generate results by attaching tools into workflows (pipelines) which can have multiple invocations, each processing different input dataset. The platform also makes integration of new tools by wrapping the tools (essentially generating tool function specific XML files) and adding them to the tool shed.

There exist offerings in the metagenomics space that use the galaxy framework, these include ASaiM (with the most end-to-end implementation of pipelines required in analysis of microbiome data)[7], MetaDEGalaxy [8], A-GAME [9], GmT [10]. All the tools provide reproducible platforms for analysing metagenomic data, however, they rely on on-device implementations or usage of Docker containers. There are webapps like EBI Metagenomics [11] and MG-RAST server [12] that provide such workflows directly on the web but have certain drawbacks like essentially being black box tools, having limited flexibility and lack of reproducibility as result of software updates and versions. Thus, there are challenges when it comes to setting up and configuring all the tools (even with Docker) and the varied execution times due to difference each software’s processing efficiency. Here we describe AutoTA, a set of workflows that use qiime2 to perform diversity analysis and taxonomic assignments, these workflows are built on a wrapper of qiime2 in the Galaxy toolshed [13]. The wrapper is available to download in the Galaxy toolshed (as of Feb 2024). We introduce these workflows as diversity analysis and taxonomic classifications are the most common initial tasks in many microbiome experiments [14](Fig 1). We provide these workflows as a starting point for building a suite of Galaxy workflows for end-to-end analysis. The qiime2 wrapper in galaxy toolshed is hosted in the toolbox on https://cancer.usegalaxy.org, which only requires the user to sign in and start using qiime right away. Our workflows are available as standalone files which can be imported into any Galaxy instance having qiime2 suite of tools. The motivation behind making workflows purely for Galaxy and using them within galaxy the way it is hosted was largely the processing challange that occur during training of classifiers especially on latest databases containing significantly large data. The latest databases like SILVA[15], GreenGenes2[16] require massive amounts of RAM to train and use classifiers. We attempted to use traditional qiime2 pipeline to train the classifier on SILVA 138.1 database on a machine running linux and having 32 GB RAM, but the process was killed (the algorithm is greedy) which makes it difficult to perform downstream steps relating to taxonomic analysis. This indicates requirement of high order RAM and processing power which are not available in every laboratory. We ran the same training process on Galaxy using the provided default backend and achieved successful completion albeit longer processing time due to the highly complex database. Therefore, workflows presented here should help scientist with limited computational resources to perform taxonomic classification and diversity analysis rapidly and reproducibly on Galaxy (cancer.usegalaxy.org).

**Figure 1.**
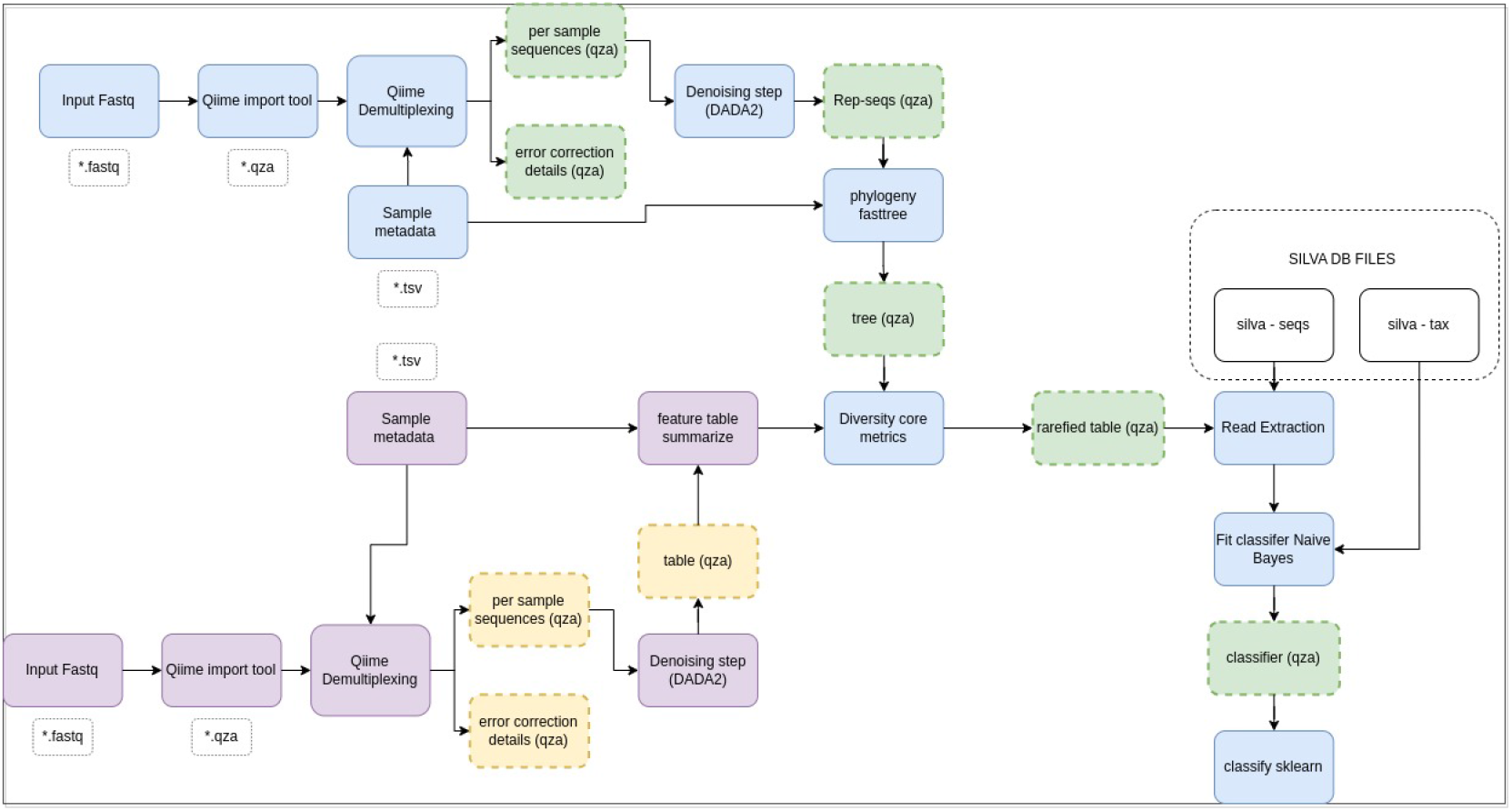
The sampling depth workflow and the main taxonomic assignment workflow is conceptually combined to give a clear picture of the analysis

**Figure 2.**
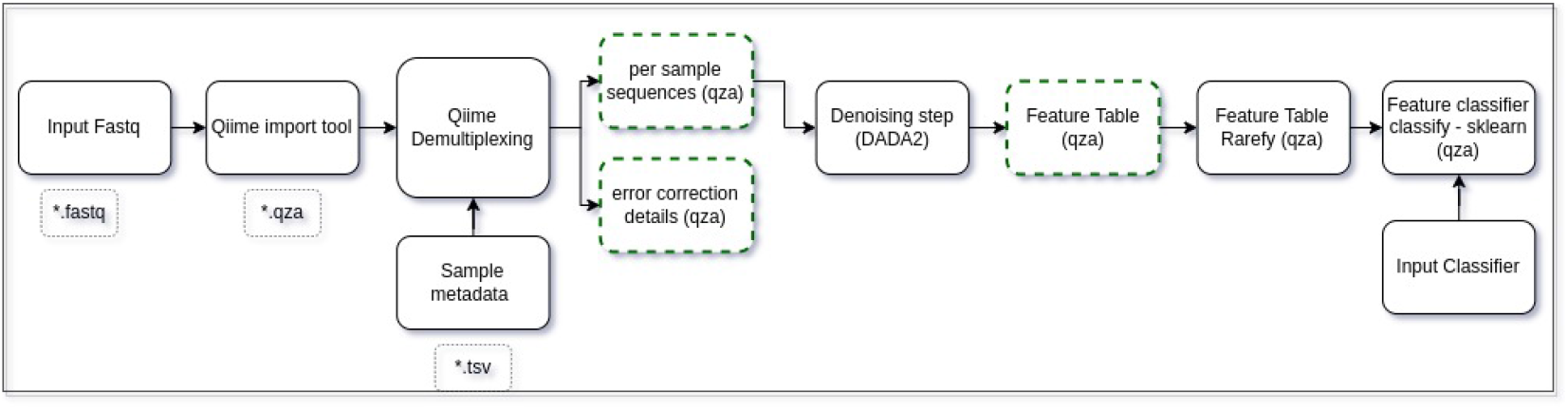
Third workflow to automate application of various classifiers (The green, dashed nodes are the generated outputs which are relevant in downstream analysis in the workflow)

**Figure 3.**
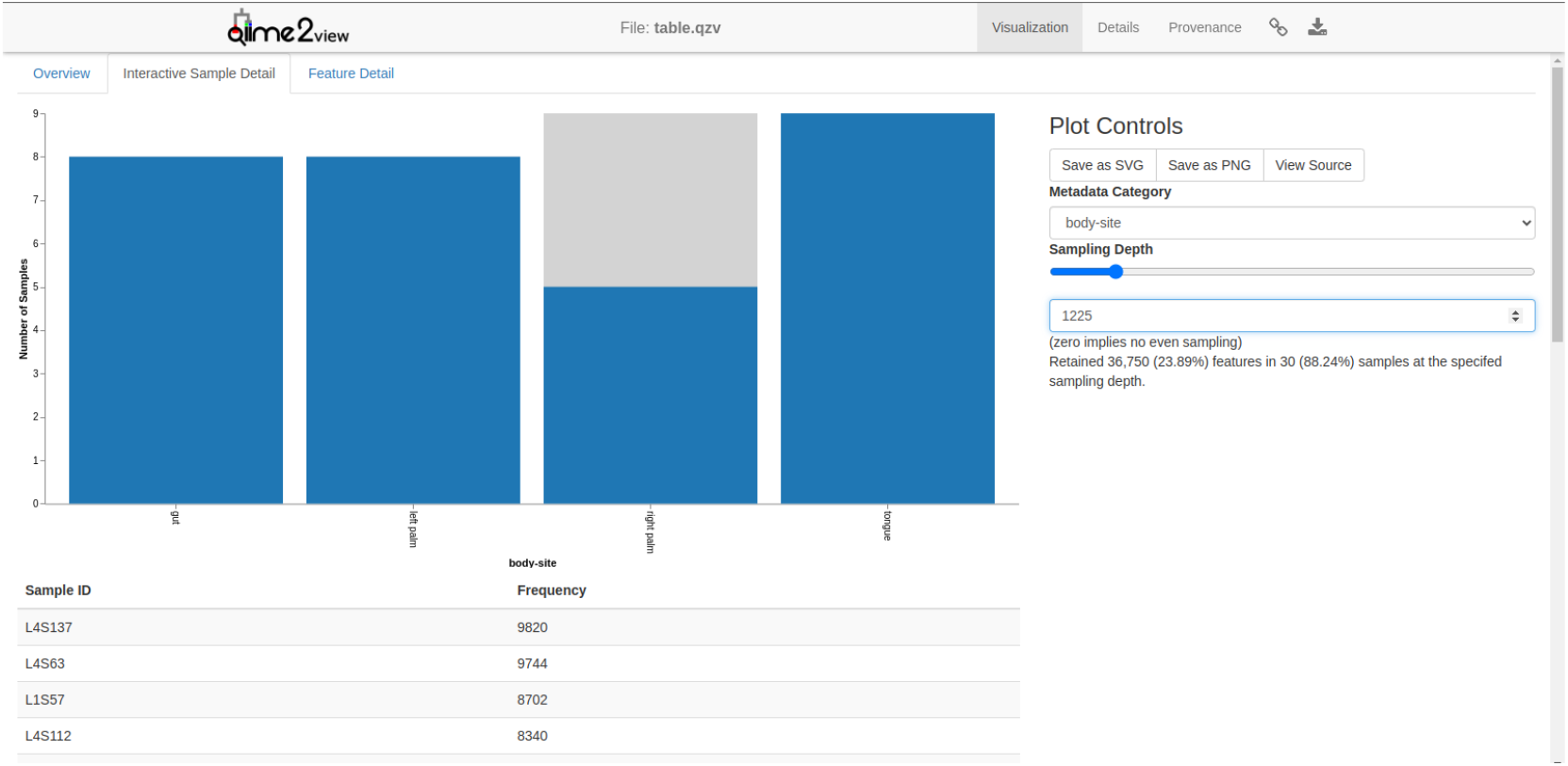
The interactive sample detail tab allows interactive determination of the sampling depth.

**Figure 4.**
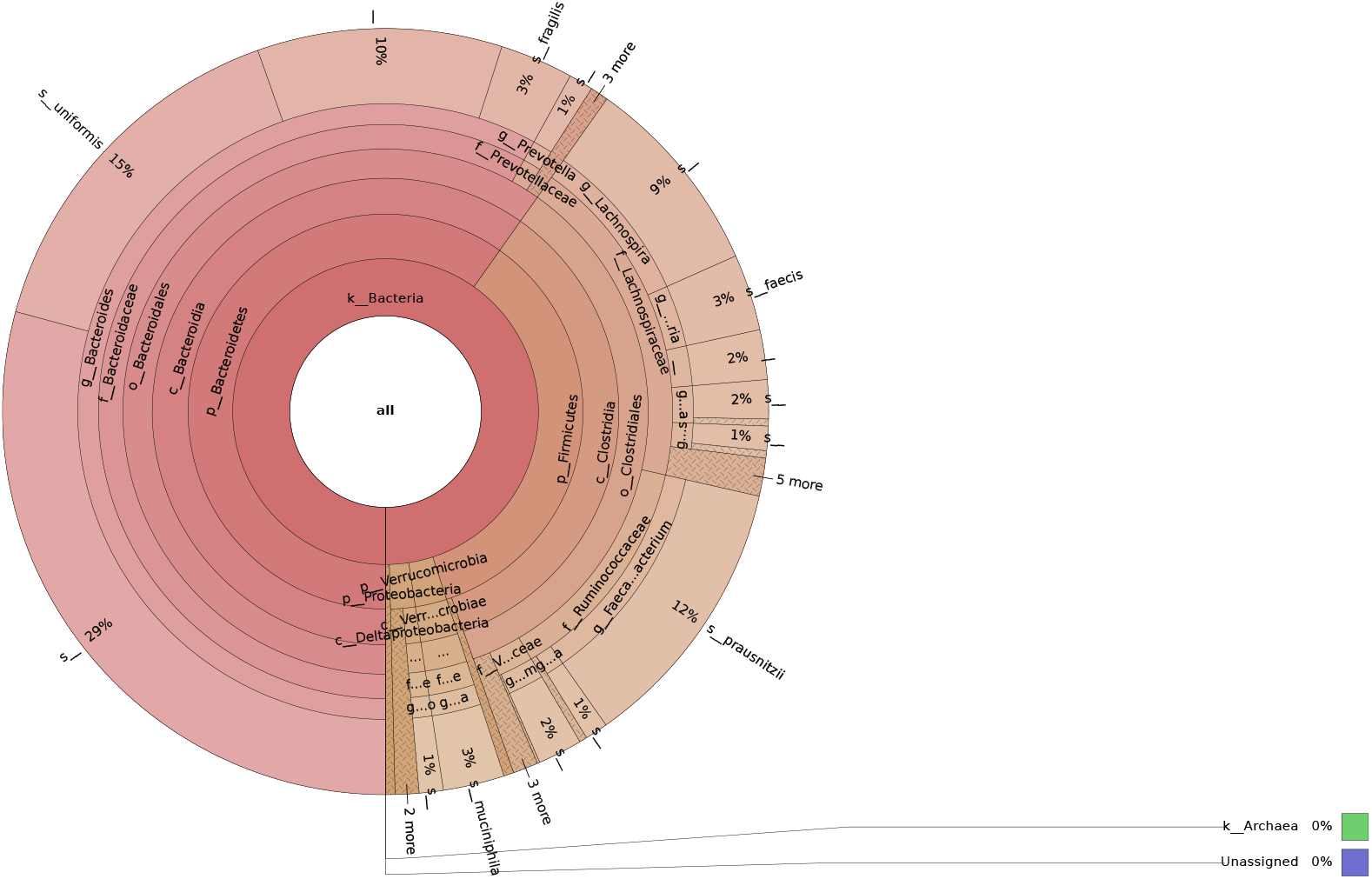
This is krona plot generated by using the taxonomy and table files mentioned in the tutorial. The classifier was a pre-trained GreenGenes classifier for V4 region.

**Figure 5.**
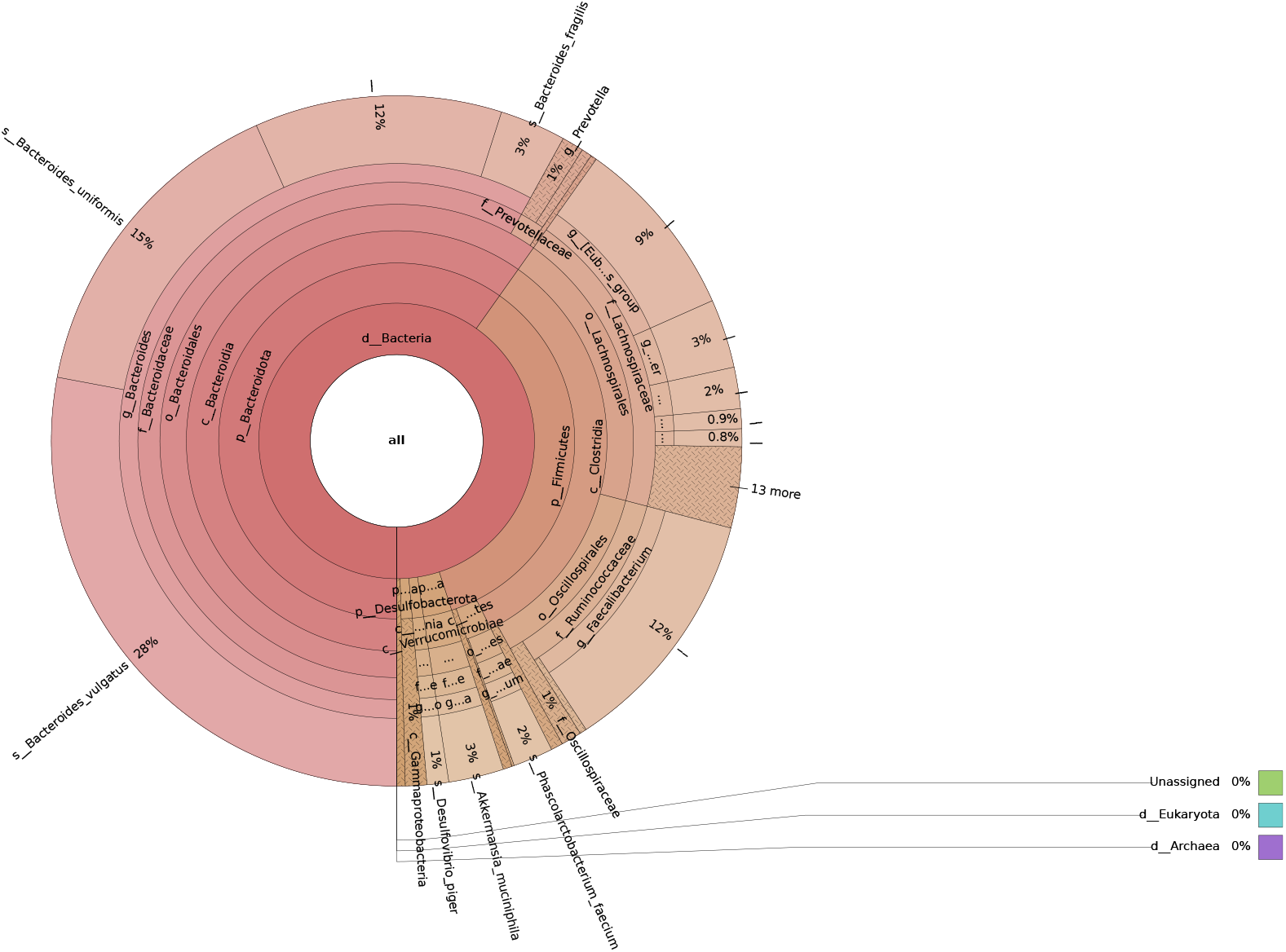
We used our workflows to generate the classifier and applied it on the tutorial input data. Our custom classifier was trained on SILVA 138.1 for V4 region.

## 2 Methods

### 2.1 Inputs

Considering that the workflows on Galaxy are completely editable and there are more than one option for each step of the analysis, most general case is presented here. For example, there is a workflow for taxonomic assignments with provisions for adding the files of silva database (sequences file and taxonomy text file), in case of the GreenGenes2, the workflow can be edited to add only one input node. The aim is to allow fastest possible way of getting started with microbiome data analysis, this could be of high value as teaching materials for novice scientists or users new to qiime2-based analysis.

The most general case is using multiplexed sequences with barcodes for each sample, hence, demultiplexing has to be performed prior to downstream analysis. The information related to samples can be stored in a metadata tab-delimited (*.tsv) file. A typical file would include columns like ‘sample-id’, ‘barcode-sequence’, ‘subject’ and associated data types for each column. Users should keep in mind that the column names are qiime specific and a list of valid column names can be found in the online documentation. This file is later required for determinatining alpha and beta diversities.

### 2.2 Workflows

We have made two workflows for taxonomic classification and diversity analysis. Due to presence of pre-trained classifiers and various database options, we have made a helper workflow to rapidly test out classifiers. This workflow uses DADA2[17] and leaves an input classifier node for the user to add the classifiers. Since the number of outputs per run of the workflow is more than 4 qza/qzv files, we recommend using separate histories for the classifier tests. The main workflow is ‘16S rRNA Analysis - Taxonomic Assignment Workflow’ which starts with diversity analysis. An important point to note here is that diversity metrics are sensitive to sampling depth which is the number of sequences in each sample, we use sampling depth for ‘rarefaction’ of our data which involves random subsampling of the sample to include only specified number of sequences. There is always a trade off between statistical power (increasing number of samples) and no of species or strains identified (increasing depth). The ‘16S rRNA Analysis - Diversity Metrics - Sampling Depth’ workflow runs the denoiser and the output table can be visualized to interactively find suitable depth. Then the sequences are rarefied which can be used by the main workflow. If the user already knows the depth, they can add it as an input in the workflow, skipping the prior step.

The third workflow is a helper workflow to test different classifiers rapidly, pre-trained on different databases or 16S rRNA gene region based classifiers trained again on different databases.

**Table 1:** Workflows and their descriptions.

| Workflows | Workflow Description |
| --- | --- |
| 16S rRNA Analysis - Diversity Metrics - Sampling Depth | This workflow is a step of the main workflow for diversity analysis. It is separated out to find the sampling depth. View the table.qzv QIIME 2 artifact, and in particular the Interactive Sample Detail tab in that visualization. Based on this, determine the sampling depth, for example, a sampling depth of 500 will subsample the counts in each table in such a way that the output table has a count of 500. |
| 16S rRNA Analysis - Taxonomic Assignment Workflow | This workflow performs diversity analysis, rarefaction and then performs taxonomic assignments. It depends on '16S rRNA Analysis - Diversity Metrics - Sampling Depth' workflow for determination of sampling depth which is required for running the core diversity metrics. |
| 16S rRNA analysis - apply classifier | This workflow is made to test various pre-trained classifiers. The main purpose of this workflow was to provide a quick way of testing different classifiers. Some classifiers are provided in qiime2 documentation. https://docs.qiime2.org/2023.9/data-resources/ (This is the latest documentation at the time of making this workflow) |

### 2.3 Operation

The advantage of using workflows within Galaxy is that they can be shared as a *.ga file, as a link, or published on the galaxy network. The file can be imported into any Galaxy instance, local or on cancer.usegalaxy.org already containing qiime2 wrapper referred to in the introduction section. Also, qiime is available on the main Galaxy website but is not actively updated and can be found in the deprecated tools section.

## 3 Results

We validated the working of our workflows on cancer.usegalaxy.org by using data and sample metadata given in qiime2 tutorial “Moving Pictures“ present in qiime docs (ver. 2023.9). We used the latest SILVA database (ver. 138.1) for training the classifier. It was found that using newer more complex databases for taxonomic classifications required computational resources which are higher than an average bioinformatics workbench (*>* 32GB RAM). In this case, using Galaxy stands to be the best route for processing larger samples with new databases and we suggest usage of our workflows on

Galaxy for rapid and efficient processing in absence of a local CPU cluster as cloud-based solutions are increasingly getting cheaper and faster [18].

The tutorial was followed till differential abundance with ANCOM section and all the output visualization artifacts were visualized using an online service view.qiime2.org.

All the steps used same parameters as mentioned in the tutorial, but, for taxonomy classification the tutorial mentions pre-trained Naive Bayes classifier trained on GreenGenes 13 8 99% OTUs database with V4 target region. We used a custom V4 classifier trained on SILVA 138.1 database. The difference in taxonomic assignments is highlighted in the krona plots generated by the krona plots plugin for qiime2 [19].

## 4 Conclusion

AutoTA is a workflow collection to streamline and simplify taxonomy classification and diversity analysis studies in large-scale metagenomics experiments. Requirement of higher compute power, number of plugins, reproducibility and command line based versions necessitated formation of these workflows. Galaxy has been used commonly by non-bioinformaticians to perform complex sets of analyses so we made the workflows on the platform itself. The workflows on galaxy made possible use of latest databases and achieve a higher degree of classification resolution. Certain drawbacks are associated with this approach, (1) the qiime2 wrapper on Galaxy is not actively maintained and updated which may give rise some issues during importing data as an artifact, (2) the user is restricted to utilize Galaxy platform and the storage limits. However, we noticed these drawbacks will be small inconveniences compared to a complete command line based approach for bench scientists. The workflows are given as ‘most usual case’ and can be modified using Galaxy’s intuitive workflow manager and editor.

